# Evolution of Single Gyroid Photonic Crystals in Bird Feathers

**DOI:** 10.1101/2020.08.27.271213

**Authors:** Vinodkumar Saranathan, Suresh Narayanan, Alec Sandy, Eric R. Dufresne, Richard O. Prum

## Abstract

Vivid, saturated structural colors are conspicuous and important features of many animals. A rich diversity of three-dimensional periodic photonic nanostructures is found in the chitinaceous exoskeletons of invertebrates. Three-dimensional photonic nanostructures have been described in bird feathers, but they are typically quasi-ordered. Here, we report bi-continuous single gyroid β-keratin and air photonic crystal networks in the feather barbs of blue-winged leafbirds *(Chloropsis cochinchinensis sensu lato)*, which have evolved from ancestral quasi-ordered channel-type nanostructures. Self-assembled avian photonic crystals may serve as inspiration for multi-functional applications, as they suggest efficient, alternative routes to single gyroid synthesis at optical length-scales, which has been experimentally elusive.

## Introduction

Many animals produce vivid, saturated structural colors via constructive light interference from diverse integumentary nanostructures with mesoscopic (~100-350 nm) long-range order or short-range translational order *(i.e.,* quasi-order) (1,2). Structural coloration is an important aspect of their appearance, and often functions in social and sexual signaling(3). Physicists and engineers are increasingly interested in animal structural coloration as a source for inspiration for mesoscale manufacture(4, 5). A remarkable diversity of structural-color producing 3D biophotonic nanostructures has been characterized within the chitinaceous exoskeletons of invertebrates(1,2, 6). While vertebrates possess one- and two-dimensional periodic photonic nanostructures(1,2), reported 3D photonic nanostructures are typically quasi-ordered, and limited to two classes of nanostructures that produce non-iridescent or isotropic structural colors in feather barbs(7).

Here, we report single gyroid photonic crystal networks of β-keratin and air within brilliantly colorful blue and green feather barbs of the Blue-winged Leafbird *(Chloropsis cochinchinensis s. l.,* Chloropseidae), revealed by synchrotron small angle X-ray scattering (SAXS) and scanning electron microscopy (SEM). We compare these ordered morphologies to homologous channel-type nanostructures with short-range order and intermediate structures from other closely-related *Chloropsis* species, and their sister group, the fairy bluebirds *(Irena,* Irenidae)(7, 8).

## Results

Light micrographs of Blue-winged Leafbird feathers show iridescent highlights within individual barb cells (Fig. 1b-d). Angle-resolved spectral measurements document weak iridescence (Fig. 1e-f). SEM images (Fig. 1g-j) reveal ordered interconnected mesoporous networks of β-keratin rods and air channels, with a polycrystalline texture. The SAXS diffraction patterns generally exhibit six-fold symmetries and up to 8 orders of discrete Bragg spots (Figs. 1k-n and 2a-c) diagnosable as single gyroid (*I4*_1_3*2*) space-group(6, 9). The observed microspectral reflection peak of a blue epaulet feather is consistent with the expected photonic bandgap structure of an appropriately-sized single gyroid (Fig. 1o).

**Figure 1.**
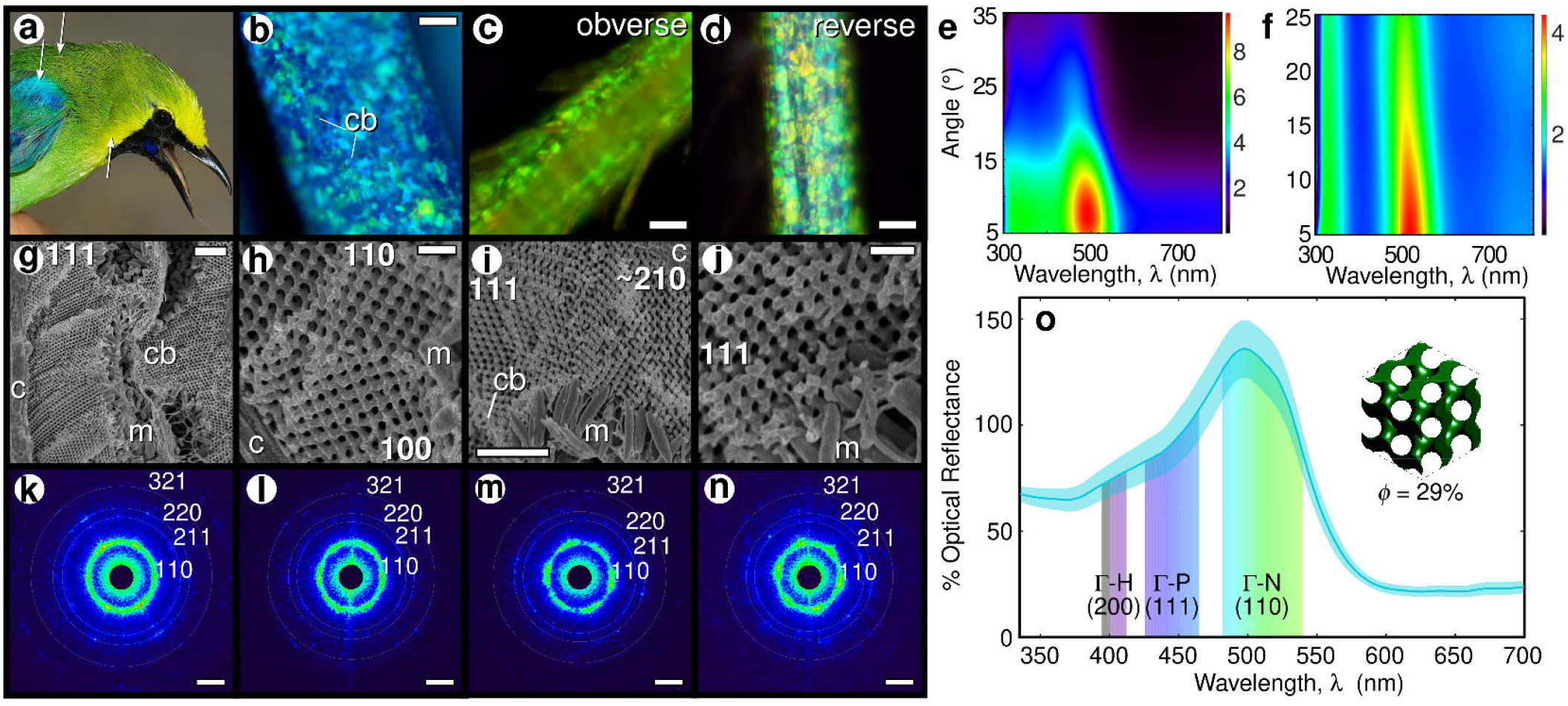
Single gyroid photonic crystals in the plumage of Blue-winged Leafbird *(Chloropsis cochinchinensis kinneari*). (**a**) Photograph with approximate sampling locations, © John C. Mittermeier. Representative light micrographs with iridescent highlights and angle-resolved specular reflectance measurements (false color) (row 1), SEM images (row 2) and SAXS diffraction patterns (row 3) of brilliant blue epaulet (**b,e,g,h,k,l**), and green back (**c,d,f,i,j,m,n**) feather barbs. Crystallite domain orientations are indicated in SEM images (**e-h**). SAXS patterns (**i-l**) are depicted in a logscale, false-color encoding and indexed as per standard IUCr conventions (white concentric circles)(6, 9). (**o**) Microspectrophotometric measurements (turquoise line – mean with std. error envelope) of blue epaulet barbs are congruent with photonic bandgap modeling (shaded pseudogaps) of a single gyroid (inset). Scale bars: **b-d** – 20 μm; **g,i** – 2 μm; **h,j** – 500 nm; **k-n** – 0.025 nm^-1^. Abbreviations: **c** – cortex, **cb** – cell boundary, **m** – melanosomes.

Coherence length (*ζ ≈ 2π/Δq*, where *Δq* is FWHM of structural correlation peaks) is a measure of crystallite domain size, and thus the extent of long-range translational order(6, 9). To investigate the evolution of single gyroids in *C. cochinchinensis,* we plot the coherence lengths of barb nanostructures from all *Chloropsis* and *Irena* species(8), in addition to known diversity of other avian *channel-type* nanostructures(7), against their peak structural correlations *(q_pk_)* (Fig. 2g). Interestingly, the diversity of barb nanostructures among *Chloropsis* species (green line) exhibit a continuum of intermediate states (green triangles), from ancestral *channel*-type nanostructures of *Irena* (blue dashed line) to the derived single gyroids of *C. cochinchinensis* (green asterisks; Fig. 2g). The quasiordered *channel*-type nanostructures within *Irena* and the two sexually monomorphic species of *Chloropsis* (*palawanensis* and *flavipennis*)(8) exhibit nearly constant peak widths (FWHM, *Δq*) relative to their dominant length-scales *(q_pk_), Q = q_pk_/Δq* ≈ 4, suggesting that scale-invariant or size-independent processes(10) underlie the assembly of quasi-ordered nanostructural states in these species. The evolution of nanostructural order from quasi-disorder within *Chloropsis* is apparent as a significant deviation from the scale-independent trend in Fig. 2g, characterized by a sharpening of the structural correlation peaks (Fig. 2a-f).

**Figure 2.**
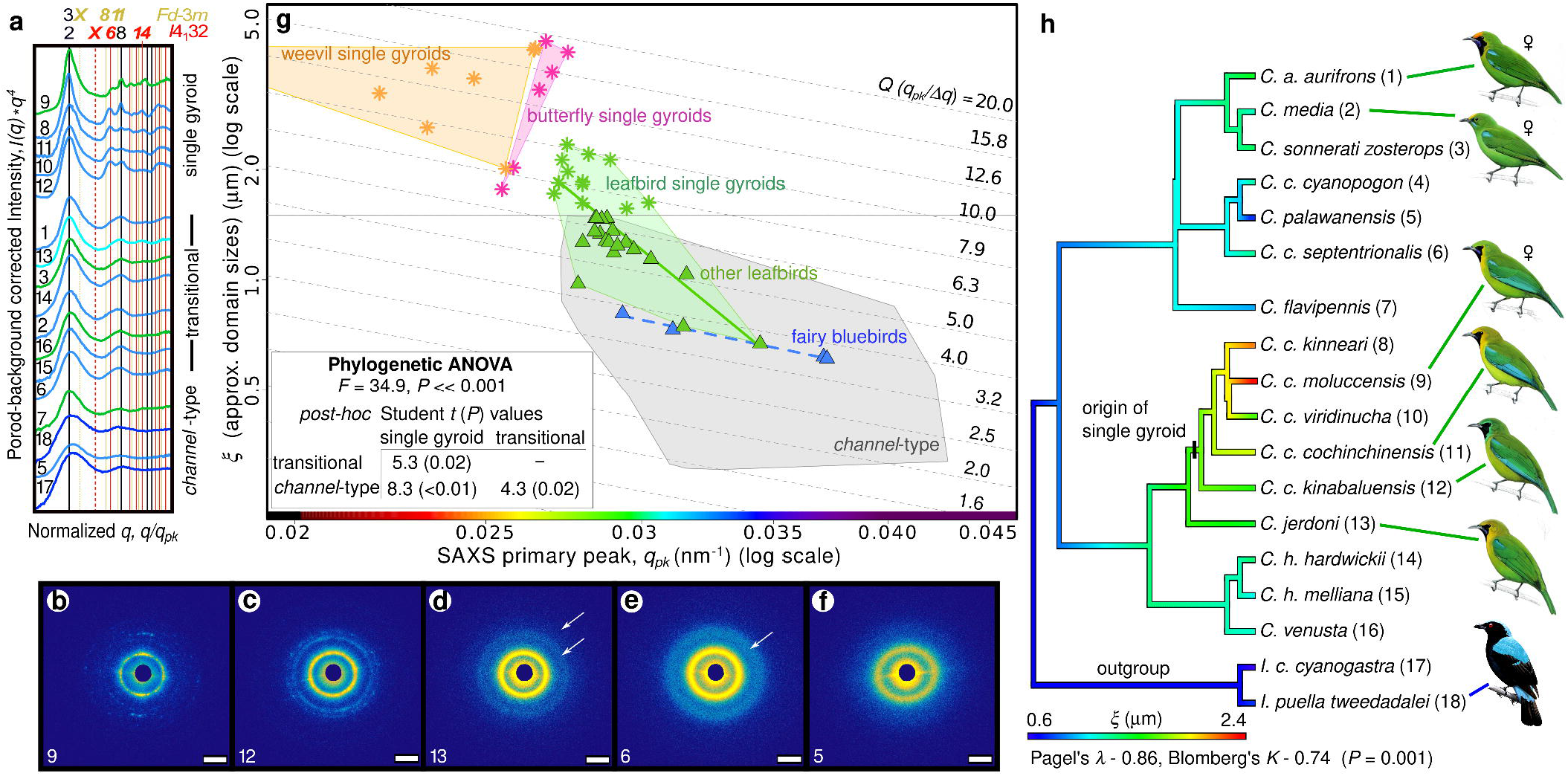
Evolutionary disorder-to-order transition in leafbird feathers. (**a**) Porod-background corrected, normalized, azimuthally-averaged SAXS profiles and (**b-f**) a representative subset of the corresponding diffraction patterns (Scale bars – 0.025 nm^-1^) of homologous feather barbs from all *Chloropsis* and 2 *Irena* species(8), in order of decreasing long-range translational order (*ζ*). Profile colors reflect the approximate hue of assayed plumage patches and numbers correspond to taxa labels in **h** (see dataset S1). White arrows in (**d,f**) indicate higher order features seen in transitional barb nanostructures. Vertical lines in (**a**) (solid – allowed, dashed – forbidden) denote expected Bragg reflections for alternative cubic space groups(6, 9). (**g**) *Ashby* diagram with nanostructural *Q*-factors *(q_pk_/Δq,* a measure of spectral purity) plotted as scale-independent (dashed gray) isolines in equally spaced deci-decades. Plotted alongside leafbird single gyroids (*), transitional and ancestral *channeltype* nanostructures of other leafbirds (green ▲) and fairy bluebirds (blue ▲) are the gamut of known avian *channel-type* nanostructures(7), and self-assembled visible length-scale single gyroids in butterflies(9) and weevils(6). Colored lines are linear regressions of log-transformed data for fairy bluebirds (dashed blue), and leafbirds (green). Shaded regions are convex hulls. The spectrum on the x-axis is an approximate color guide to barb hues, for a given *q_pk_*. Inset shows the results of phylogenetic ANOVA. (**h**) Bayesian ancestral state reconstruction of *ζ* on a consensus molecular phylogeny of *Chloropsis* and *Irena*(8), with a single origin of single gyroid marked in the blue-winged leafbird clade *(Chloropsis cochinchinensis s. l.).* Illustrations: *Chloropsis spp*. – John Gale/Naturalis Leiden (CC-BY 4.0), and *Irena puella* – © Lynx Edicions.

Because the plumage photonic nanostructures of *Chloropsis* and *Irena* are homologs that first evolved in the most recent common ancestor of these two monophyletic sister genera(7, 8), our comparative analysis demonstrates that the single gyroid photonic networks of blue-winged leafbirds were evolutionarily derived from ancestral quasi-ordered *channel-type* networks, still present in *Irena* (Fig. 2h). Moreover, the variation in nanostructures among *Chloropsis* species indicates the evolutionary transition from disorder-to-order proceeds as a gradual appearance and sharpening of higher-order Bragg reflections(*cf*. 6, 9). In addition to a second-order feature that is typical of *channel-type* nanostructures(7), the azimuthal SAXS profiles of the transitional *Chloropsis* nanostructures exhibit 1 to 2 additional higher-order peaks at ratios close to √6-√8, √14, and √22-√24, but not √4 (Fig. 2a).

## Discussion

The origin of 3D photonic crystals from quasi-ordered nanostructures in blue-winged leafbirds are evolutionarily parallel with the derivation of 2D hexagonal columnar crystals in *Philepitta* from ancestral 2D quasi-ordered arrays of parallel collagen fibers in structurally colored skin of the most recent common ancestor with *Neodrepanis* (Philepittidae)(11). In both cases, evolutionary transitions from quasi-ordered to ordered photonic crystals with much narrower structural correlation peaks (Fig. 2a-f, and Figs. 6 and 7 of (11)) result in the production of highly saturated or purer hues that are readily perceivable by avian visual systems(3). These results suggest social or sexual selection on perceivable optical properties of biophotonic nanostructures *(e.g.,* preferences for purer/ more saturated hues) have likely driven evolutionary transitions in nanostructural spatial organization, resulting in the evolution of extraordinarily brilliant structural coloration(11).

Single gyroids that can exhibit large complete bandgaps have long been a target for photonic and photovoltaic engineering (reviewed in 5), but synthetic self-assembly of single gyroids at visible length-scales has proven elusive. In insect wing scales, single gyroid photonic crystals are templated by the co-option of the innate ability of cell membranes to invaginate into core-shell double-gyroid precursor networks(6, 9). The chitinous cuticle polymerizes within extracellular space enclosed by the plasma membrane, leaving behind a single gyroid network of chitin in air, upon apoptosis. Current synthetic approaches to self-assemble single gyroids follow a similar symmetry-breaking pathway (reviewed in 5), starting with a double gyroid or an alternating gyroid in a di- or tri-block copolymer and subsequent selective etching of the matrix and one of the two network phases. Synthetic single gyroids produced this way are limited to small lattice parameters (typically < 100 nm). Synthetic lipid water systems also show cubic phases including double gyroid but with lattice parameters limited to just tens of nm(12).

Quasi-ordered *channel-type* photonic nanostructures of avian feather barbs are understood to self-assemble within medullary cells via self-arrested phase separation of polymerizing β-keratin from the cytoplasm, in the absence of any cytoskeletal prepatterns or membraneous precursor templates (reviewed in 13). Our findings suggest that single gyroid photonic crystals may be efficiently synthesized at optical length-scales through processes akin to bottom-up phase-separation. Simulations(14) suggest that the phase behavior of patchy particles with mutual short-range attraction and long-range repulsion can include a metastable single gyroid phase. Future research into *in vivo* and *in vitro* arrested phase separation of colloidal solutions of charged proteins or similar polymers may provide novel insights into synthetic self-assembly of ordered mesoporous phases, including the single gyroid(4).

## Materials and Methods

See extended methods provided in SI Appendix.

## Acknowledgments

We thank ornithology curators and staff of Yale Peabody Museum, and Lee Kong Chian Museum of Natural History for loans and access to study-skins. Michael Rooks assisted with SEM. V.S. is grateful to Dan Morse and UCSB Institute for Collaborative Biotechnologies for support during a sabbatical that enabled some SAXS data acquisition.

## Data Availability

All study data are included in the article, and dataset S1.

## References

1. Vukusic P & Sambles JR (2003) Photonic structures in biology. Nature 424(6950):852–855.

2. Kinoshita S, Yoshioka S, & Miyazaki J (2008) Physics of structural colors. Reports on Progress in Physics 71:076401.

3. Cuthill IC, et al. (2017) The biology of color. Science 357(6350).

4. McDougal A, Miller B, Singh M, & Kolle M (2019) Biological growth and synthetic fabrication of structurally colored materials. Journal of Optics 21 (7):073001.

5. Dolan JA, et al. (2015) Optical Properties of Gyroid Structured Materials: From Photonic Crystals to Metamaterials. Advanced Optical Materials 3(1):12–32.

6. Saranathan V, et al. (2015) Structural Diversity of Arthropod Biophotonic Nanostructures Spans Amphiphilic Phase-Space. Nano Lett 15(6):3735–3742.

7. Saranathan V, et al. (2012) Structure and Optical Function of Amorphous Photonic Nanostructures from Avian Feather Barbs: A Comparative Small Angle X-ray Scattering (SAXS) Analysis of 230 Bird Species. Journal of The Royal Society Interface 9:2563–2580.

8. Moltesen M, Irestedt M, Fjeldsa J, Ericson PG, & Jonsson KA (2012) Molecular phylogeny of Chloropseidae and Irenidae – cryptic species and biogeography. Mol Phylogenet Evol 65(3):903–914.

9. Saranathan V, et al. (2010) Structure, function, and self-assembly of single network gyroid (I4(1)32) photonic crystals in butterfly wing scales. P Natl Acad Sci Usa 107(26):11676–11681.

10. Fratzl P, Lebowitz JL, Penrose O, & Amar J (1991) Scaling functions, self-similarity, and the morphology of phase-separating systems. Physical Review B 44(10):4794–4811.

11. Prum RO, Torres RH, Kovach C, Williamson S, & Goodman SM (1999) Coherent light scattering by nanostructured collagen arrays in the caruncles of the Malagasy asities (Eurylaimidae: Aves). Journal of Experimental Biology 202:3507–3522.

12. Luzzati V & Spegt PA (1967) Polymorphism of Lipids. Nature 215(5102):701–704.

13. Saranathan V & Finet C (2020) Cellular and Developmental Basis of Avian Structural Coloration. arXiv 2012.10338.

14. Stopper D & Roth R (2017) Phase behavior and bulk structural properties of a microphase former with anisotropic competing interactions: A density functional theory study. Phys Rev E 96(4):042607.

