## Supplementary Information for "Evolution of Single Gyroid Photonic Crystals in Bird Feathers"

**This file includes:**

Supplementary text

Supplementary Dataset 1

References for SI

**Extended Methods**

***Specimens:*** The nanostructure and structural color production in feather barbs of all 10-15 putative *Chloropsis* (Chloropseidae) and 2 *Irena* (Irenidae) species(1) were analyzed in this study (see Dataset S1).

***Synchrotron Small Angle X-ray Scattering (SAXS):*** In order to average over as few barb cells as possible, pinhole SAXS at either 15x15 or 10x10 μm (horizontal x vertical) sizes was performed in transmission geometry on feather barbs at beamline 8-ID-I of the Advanced Photon Source (Argonne National Labs) and acquired data was processed as per standard protocol(2-4).

***Scanning Electron Microscopy (SEM):*** For scanning electron microscopy (SEM), fractured barb samples were gold- or platinum-coated and imaged on a Hitachi SU-70 environmental SEM.

***Angle-resolved spectrophotometry:*** Angle-resolved UV-VIS-NIR reflectance spectra (2 s integration time, 5x averaging) from individual feathers were measured in specular ($\theta$-$2\theta$) geometry (5) using a custom goniometer setup coupled to an Ocean Optics USB2000+ spectrophotometer and an Ocean Optics DH-BAL 2000 light source, as described elsewhere(2). Reflectance was calibrated using an Ocean Optics Spectralon matte white standard and a piece of matt-black velvet cloth for dark reference.

***Normal incidence spectrophotometry:*** UV-VIS-NIR normal incidence reflectance measurements (0.1 – 0.4 s integration time) were acquired from a ~5 $mm^2$ illuminated patch, using a bifurcated probe and holder either from individual feathers (with 3x averaging) or study skins (2-3 different locations per plumage patch), as per standard protocol(2).

***UV-VIS-NIR Microspectropectrophotometry and Light Microscopy:*** Normal-incidence UV-VIS-NIR reflectance microspectra of leafbird feather barbs were acquired using a uSight-2000-Ni microspectrophotometer (Technospex Pte. Ltd., Singapore). Spectra with usable range between 335-950 nm were collected using a high NA 100x objective from a ~1.5 μm sized spot (100 ms integration time, 40x averaging) and calibrated using an Avantes WS-2 matt-white standard. Light microscope images were recorded using the same microspectrometer setup at various magnifications, using a Touptek U3CMOS-05 camera.

***Photonic Bandgap modeling:*** The MIT photonic bandgap package (MPB)(6) was used to calculate the first eight bands of a single gyroid photonic crystal, using a mesh-size of 5 and a resolution of 32 to discretize the unit cell. As in (4), the bandgap calculations were optimized to find the dielectric (βkeratin, refractive index = 1.58) volume fractions needed ($\phi$ = 0.29) to produce a mid-gap frequency (0.6043) of the Γ-N (110) pseudgogap, as given by $a / \lambda_{pk}$, where $a$ is the lattice parameter measured using SAXS and $\lambda_{pk}$ is the microspectral peak hue of the blue epaulet of *C. cochinchinensis kinneari* (see Fig. 1o, and Dataset S1).

***Phylogenetic analyses:*** Bayesian continuous ancestral character state reconstructions of coherence lengths ($\xi$) were performed in *R* (version 3.6), using the *anc.Bayes()* routine of the *phytools* package (version 0.7-47)(7), with 1,000,000 generations (20% burn-in). The average coherence length of the two *Irena* species was set as the informative prior for the root/ ancestral node of *Irena* and *Chloropsis*. Using the *contMap()* function, the ancestral state reconstructions were mapped onto the topology, which was the 50% majority rule consensus molecular phylogeny of *Irena* and *Chloropsis* presented in Fig. 6 of (1). Polytomies (placement of *C. flavipennis*, and the "*C. moluccensis*" clade) (1) were randomly resolved and zero branch lengths were set to 1/100 of the total tree length. Phylogenetic signal (Blomberg's *K* and Pagel's $\lambda$) were computed using the *phylosig()* routine. Phylogenetically-corrected analyses of variance (n = 1000 simulations) and *post-hoc* tests were performed using the *phylANOVA()* routine of *phytools*(7).

**Dataset S1**

A spreadsheet detailing bird study-skin sampling information as well as the optical reflectance and nanostructural parameters of all plumage patches of leafbirds and fairy bluebirds assayed in this study. For details of other avian *channel*-type barb nanostructures plotted in Fig. 2, see Tables S1 and S2 of (2); for butterfly and weevil single gyroids, see Table S1 of (4) and Table S1 of (3) respectively.

**Supplementary References**


1. Moltesen M, Irestedt M, Fjeldsa J, Ericson PG, & Jonsson KA (2012) Molecular phylogeny of Chloropseidae and Irenidae - cryptic species and biogeography. *Mol Phylogenet Evol* 65(3):903-914.
2. Saranathan V, *et al.* (2012) Structure and Optical Function of Amorphous Photonic Nanostructures from Avian Feather Barbs: A Comparative Small Angle X-ray Scattering (SAXS) Analysis of 230 Bird Species. *Journal of The Royal Society Interface* 9:2563–2580.
3. Saranathan V, *et al.* (2015) Structural Diversity of Arthropod Biophotonic Nanostructures Spans Amphiphilic Phase-Space. *Nano Lett* 15(6):3735-3742.

4. Saranathan V, *et al.* (2010) Structure, function, and self-assembly of single network gyroid (I4(1)32) photonic crystals in butterfly wing scales. *P Natl Acad Sci Usa* 107(26):11676-11681.
5. Noh H, *et al.* (2010) How Noniridescent Colors Are Generated by Quasi-ordered Structures of Bird Feathers. *Advanced Materials* 22(26-27):2871-2880.
6. Johnson S & Joannopoulos J (2001) Block-iterative frequency-domain methods for Maxwell's equations in a planewave basis. *Opt. Express* 8(3):173-190.
7. Revell LJ (2012) phytools: an R package for phylogenetic comparative biology (and other things). *Methods in Ecology and Evolution* 3(2):217-223.
